# Host adaptation and convergent evolution increases antibiotic resistance without loss of virulence in a major human pathogen

**DOI:** 10.1101/370940

**Authors:** Alicia Fajardo-Lubián, Nouri L. Ben Zakour, Alex Agyekum, Jonathan R. Iredell

## Abstract

As human population density and antibiotic exposure increases, specialised bacterial subtypes have begun to emerge. Arising among species that are common commensals and infrequent pathogens, antibiotic-resistant ‘high-risk clones’ have evolved to better survive in the modern human. Here, we show that the major matrix porin (OmpK35) of *Klebsiella pneumoniae* is not required in the mammalian host for colonisation, pathogenesis, nor for antibiotic resistance, and that it is commonly absent in pathogenic isolates. This is found in association with, but apparently independent of, a highly specific change in the co-regulated partner porin, the osmoporin (OmpK36), which provides enhanced antibiotic resistance without significant loss of fitness in the mammalian host. These features are common in well-described ‘high-risk clones’ of *K. pneumoniae*, as well as in unrelated members of this species and similar adaptations are found in other members of the Enterobacteriaceae that share this lifestyle. Available sequence data indicates evolutionary convergence, with implications for the spread of lethal antibiotic-resistant pathogens in humans.

**Author summary:** *Klebsiella pneumoniae* is a Gram-negative enterobacteria and a significant cause of human disease. It is a frequent agent of pneumonia, and systemic infections can have high mortality rates (60%). OmpK35 and OmpK36 are the major co-regulated outer membrane porins of *K. pneumoniae*. OmpK36 absence has been related to antibiotic resistance but decreased bacterial fitness and diminished virulence. A mutation that constricts the porin channel (Gly134Asp135 duplication in loop 3 of the porin, OmpK36GD) has been previously observed and suggested as a solution to the fitness cost imposed by loss of OmpK36.

In the present study we constructed isogenic mutants to verify this and test the impact of these porin changes on antimicrobial resistance, fitness and virulence. Our results show that loss of OmpK35 has no significant cost in bacterial survival in nutrient-rich environments nor in the mammalian host, consistent with a predicted role outside that niche. When directly compared with the complete loss of the partner osmoporin OmpK36, we found that isogenic OmpK36GD strains maintain high levels of antibiotic resistance and that the GD duplication significantly reduces neither gut colonisation nor pathogenicity in a pneumonia mouse model. These changes are widespread in unrelated genomes. Our data provide clear evidences that specific variations in the loop 3 of OmpK36 and the absence of OmpK35 in *K. pneumoniae* clinical isolates are examples of successful adaptation to human colonization/infection and antibiotic pressure, and are features of a fundamental evolutionary shift in this important human pathogen.

## Introduction

Host adaptation and niche specialisation is well described in bacteria. As human population density rises, commensals and pathogens among the Enterobacteriaceae are transmitted directly from human to human and increasingly exposed to antibiotics. *K. pneumoniae* is now a common cause of healthcare-associated infections and is one of the most important agents of human sepsis [1]. High morbidity and mortality is associated with acquired antibiotic resistance, most importantly by horizontal transfer of genes encoding extended-spectrum β-lactamases (ESBL) [2] and plasmid-mediated AmpC β-lactamases (pAmpC) [3]. Carbapenem antibiotics have been effective against such isolates for decades but resistance to these antibiotics is increasingly common in turn [4] and in February 2017, carbapenem resistant Enterobacteriaceae were listed as highest (‘critical’) research priorities by the World Health Organisation. Acquired genes encoding efficient carbapenem hydrolysing enzymes [5] typically require phenotypic augmentation by permeability reduction to be clinically relevant in the Enterobacteriaceae. Indeed, clinically significant carbapenem resistance may even be seen with the less specialised AmpC or ESBL enzymes in strains with sufficiently reduced outer membrane permeability [6, 7].

*K. pneumoniae* expresses two major nonspecific porins (OmpK35 and OmpK36) through which nutrients and other hydrophilic molecules such as β-lactams diffuse into the cell [8, 9]. The expression of these two major porins in *K. pneumoniae* are strongly linked with β-lactam susceptibility [6, 7] and strains lacking both porins exhibit high levels of resistance [10]. *K. pneumoniae* is commonly present in the human gut [1] but also grows in low-nutrient and low-osmolarity conditions, with decreased expression of the ‘osmoporin’, OmpK36, and increased expression of the ‘matrix porin’, OmpK35, which has the greater general permeability. In the mammalian host *in vivo*, and in nutritious media *in vitro*, OmpK36 is the principal general porin and the gateway for β-lactam antibiotics, these being the most frequently prescribed antibiotic class in humans and the cornerstone of therapy for serious infections.

The fitness cost of certain antibiotic resistance mutations is well described [11-14]. Significantly reduced expression of porins provides some protection from β-lactam antibiotics but may incur a considerable metabolic cost as vital nutrients are simultaneously excluded [15]. Outer membrane permeability is thus a balance between self-defence and competitive fitness [16, 17]. Global antibiotic restriction policies are founded on the premise of an inverse relationship between competitive fitness and resistance to antibiotics [18] and the expectation that antibiotic-resistant mutants will fail to successfully compete with their antibiotic-susceptible ancestors [19]. However, analysis of the principal porin relevant to infection in the mammalian host, OmpK36, revealed a key role for a transmembrane β-strand loop (loop3, L3) in the porin inner channel (‘eyelet’), which is electronegative at physiological pH. Minor changes in this region have been observed that are expected to be relatively permissive of small nutrient molecule diffusion but which may exclude more bulky anionic carbapenem and cephalosporin antibiotics [20].

Highly antibiotic-resistant *K. pneumoniae* is both a critical threat pathogen and a model of adaptation in a world with increasing human density and antibiotic exposure. The aim of this study was therefore to understand the pathogenesis and antimicrobial resistance implications of common changes in major porins that diminish membrane permeability.

## Materials and methods

### Bacterial strains, plasmids, primers and growth conditions

Bacterial strains, plasmids and primers used in this study are listed in Tables 1 and S1. Porin mutants were constructed in three antibiotic-susceptible *K. pneumoniae* strains (ATCC 13883, and clinical isolates 10.85 and 11.76 from our laboratory). Bacterial isolates were stored at −80°C in Nutrient broth (NB) with 20% glycerol and recovered on LB agar plates. Unless otherwise indicated, strains were routinely grown in Mueller-Hinton broth (MHB, BD Diagnostics, Franklin lakes, NJ, USA) or Luria-Bertani (LB, Life Technologies, Carlsbad, CA, USA). *E. coli* and *K. pneumoniae* strains carrying the chloramphenicol-resistant plasmids pKM200 and pCACtus were grown at 30°C on LB agar or in LB broth supplemented with 20 μg/ml chloramphenicol (Sigma-Aldrich, St. Louis, MO, USA). The growth of bacterial cells was determined by measuring the optical density at 600 nm (OD_600_) in an Eppendorf Biophotometer (Eppendorf AG, Hamburg, Germany).

**Table 1.**
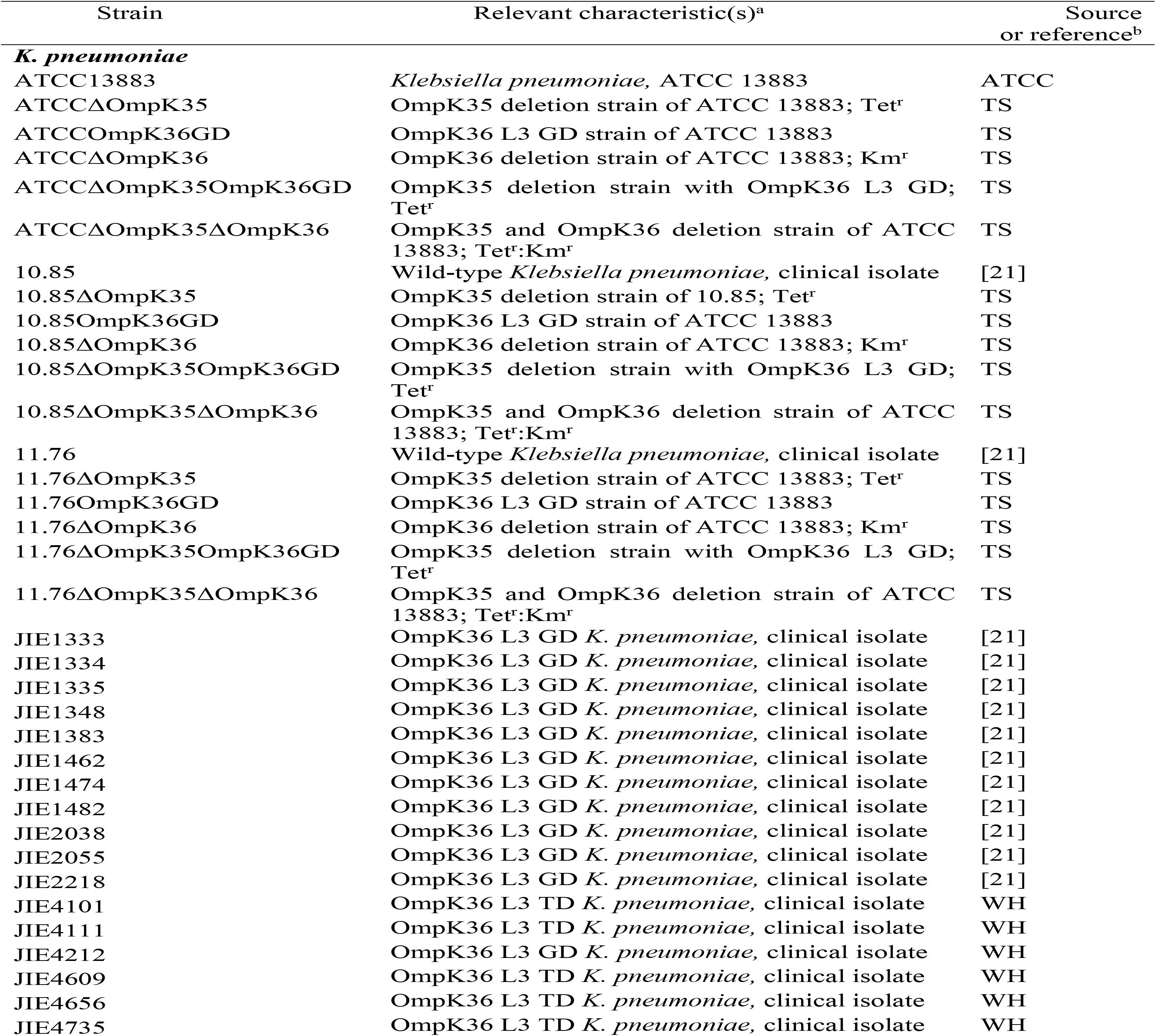
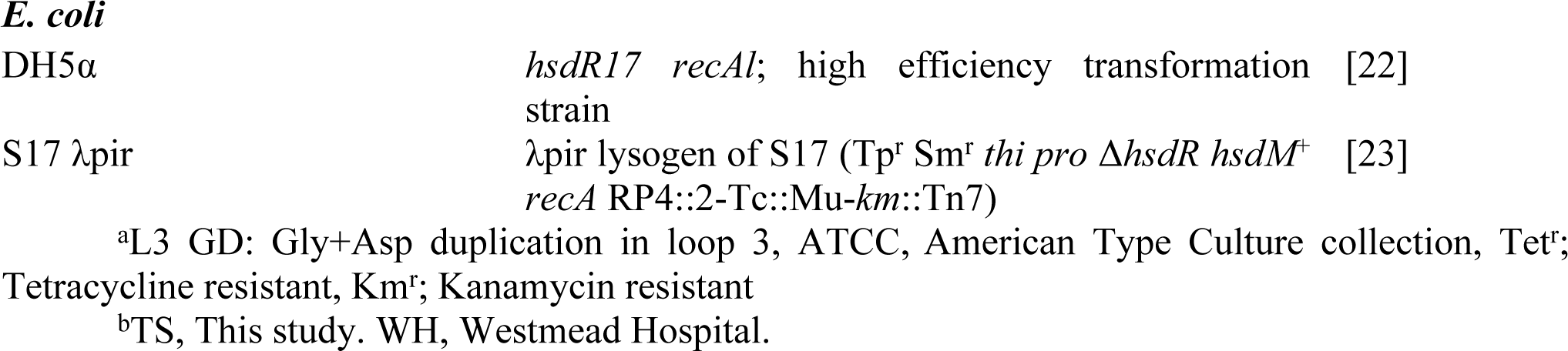
Bacterial strains used in this study

| Strain | Relevant characteristic(s)<sup>a</sup> | Source or reference<sup>b</sup> |
| --- | --- | --- |
| ***K. pneumoniae*** |  |  |
| ATCC13883 | *Klebsiella pneumoniae*, ATCC 13883 | ATCC |
| ATCCΔOmpK35 | OmpK35 deletion strain of ATCC 13883; Tet<sup>r</sup> | TS |
| ATCCOmpK36GD | OmpK36 L3 GD strain of ATCC 13883 | TS |
| ATCCΔOmpK36 | OmpK36 deletion strain of ATCC 13883; Km<sup>r</sup> | TS |
| ATCCΔOmpK35OmpK36GD | OmpK35 deletion strain with OmpK36 L3 GD; Tet<sup>r</sup> | TS |
| ATCCΔOmpK35ΔOmpK36 | OmpK35 and OmpK36 deletion strain of ATCC 13883; Tet<sup>r</sup>:Km<sup>r</sup> | TS |
| 10.85 | Wild-type *Klebsiella pneumoniae*, clinical isolate | [21] |
| 10.85ΔOmpK35 | OmpK35 deletion strain of 10.85; Tet<sup>r</sup> | TS |
| 10.85OmpK36GD | OmpK36 L3 GD strain of ATCC 13883 | TS |
| 10.85ΔOmpK36 | OmpK36 deletion strain of ATCC 13883; Km<sup>r</sup> | TS |
| 10.85ΔOmpK35OmpK36GD | OmpK35 deletion strain with OmpK36 L3 GD; Tet<sup>r</sup> | TS |
| 10.85ΔOmpK35ΔOmpK36 | OmpK35 and OmpK36 deletion strain of ATCC 13883; Tet<sup>r</sup>:Km<sup>r</sup> | TS |
| 11.76 | Wild-type *Klebsiella pneumoniae*, clinical isolate | [21] |
| 11.76ΔOmpK35 | OmpK35 deletion strain of ATCC 13883; Tet<sup>r</sup> | TS |
| 11.76OmpK36GD | OmpK36 L3 GD strain of ATCC 13883 | TS |
| 11.76ΔOmpK36 | OmpK36 deletion strain of ATCC 13883; Km<sup>r</sup> | TS |
| 11.76ΔOmpK35OmpK36GD | OmpK35 deletion strain with OmpK36 L3 GD; Tet<sup>r</sup> | TS |
| 11.76ΔOmpK35ΔOmpK36 | OmpK35 and OmpK36 deletion strain of ATCC 13883; Tet<sup>r</sup>:Km<sup>r</sup> | TS |
| JIE1333 | OmpK36 L3 GD *K. pneumoniae*, clinical isolate | [21] |
| JIE1334 | OmpK36 L3 GD *K. pneumoniae*, clinical isolate | [21] |
| JIE1335 | OmpK36 L3 GD *K. pneumoniae*, clinical isolate | [21] |
| JIE1348 | OmpK36 L3 GD *K. pneumoniae*, clinical isolate | [21] |
| JIE1383 | OmpK36 L3 GD *K. pneumoniae*, clinical isolate | [21] |
| JIE1462 | OmpK36 L3 GD *K. pneumoniae*, clinical isolate | [21] |
| JIE1474 | OmpK36 L3 GD *K. pneumoniae*, clinical isolate | [21] |
| JIE1482 | OmpK36 L3 GD *K. pneumoniae*, clinical isolate | [21] |
| JIE2038 | OmpK36 L3 GD *K. pneumoniae*, clinical isolate | [21] |
| JIE2055 | OmpK36 L3 GD *K. pneumoniae*, clinical isolate | [21] |
| JIE2218 | OmpK36 L3 GD *K. pneumoniae*, clinical isolate | [21] |
| JIE4101 | OmpK36 L3 TD *K. pneumoniae*, clinical isolate | WH |
| JIE4111 | OmpK36 L3 TD *K. pneumoniae*, clinical isolate | WH |
| JIE4212 | OmpK36 L3 GD *K. pneumoniae*, clinical isolate | WH |
| JIE4609 | OmpK36 L3 TD *K. pneumoniae*, clinical isolate | WH |
| JIE4656 | OmpK36 L3 TD *K. pneumoniae*, clinical isolate | WH |
| JIE4735 | OmpK36 L3 TD *K. pneumoniae*, clinical isolate | WH |

### Construction of porin mutants

Chemical transformation, conjugations and electroporation were carried out using standard protocols. Platinum pfx DNA polymerase (Invitrogen, USA) was used to amplify blunt-ended PCR products. All PCR products were purified (PureLink Quick PCR Purification Kit; Invitrogen, USA). PCR and sequencing was used to confirm all constructs. Genomic DNA extractions were performed using a DNeasy Blood and Tissue kit (Qiagen, Valencia, CA, USA) and plasmid DNA using a PureLink Quick Plasmid Miniprep kit (Life Technologies, Carlsbad, CA, USA) or a HiSpeed Plasmid Midi Kit (Qiagen, Valencia, CA, USA).

Porin deletions mutants of *K. pneumoniae* ATCC 13883, 10.85 and 11.76 were created by introduction of *tetA* (tetracycline-resistance) or *aphA-3* (kanamycin-resistance) into unique sites in *ompK35* and *ompK36* (HincII and StuI, respectively) which had been previously cloned into pGEM-T easy (Promega, Madison, WI, USA). The disrupted porin genes were then cloned into pCACtus suicide temperature-sensitive vector (pJIAF-7 to PJIAF-12) to replace the respective chromosomal genes by homologous recombination [24]. Confirmation of correct single-copy chromosomal mutations were finally verified by PCR (Table S1).

OmpK36GD mutants were obtained by amplification of OmpK36 from each parental strain using K36GD1 / K36GD2 and K36GD3 / K36GD4 primers (Table S1). The amplicon, containing a GD duplication in L3, was cloned first in pGEM-T easy and after digestion with *Sph1* and *Sac1* (New England Biolabs, MA, USA) was introduced into pCACtus. The pCACtus-based construct (pJIAF-13 to pJIAF-18) was transformed into S17λpir and conjugated into *K. pneumoniae* ∆OmpK36 (kanamycin-resistance mutant) where the interrupted gene was replaced by OmpK36 porin with GD duplication in L3 by homologous recombination. Mutants were selected by loss of kanamycin resistance and confirmed by PCR and sequencing.

Double mutants (∆OmpK35∆OmpK36 and ∆OmpK35OmpK36GD) were constructed using lambda Red-mediated recombineering as described previously [25, 26], with some modifications. A tetracycline cassette flanked by OmpK35 deletion (~2.5 kb in size) was PCR amplified from an OmpK35 deletion mutant (∆OmpK35; tetracycline resistant-previously obtained) using primers ompK35X-F and ompK35X-R (Table S1) and PCR products were purified. Red Helper plasmid pKM200 was electroporated into ∆OmpK36 or OmpK36GD single mutants. *ompK35:tetA* fragments were electroporated into ∆OmpK36 or OmpK36GD clones carrying pKM200. Bacteria were grown at 30°C for 2 h with agitation (225 rpm) followed by overnight incubation at 37°C. Different dilutions of the electroporated cells were spread on LB agar plates containing 10 μg/ml tetracycline to select for transformants at 37°C. The correct structure was confirmed by sequencing of PCR amplicons (primers ompK35F1 and ompK35R2, Table S1).

### Antimicrobial susceptibility tests

Susceptibilities to cefazolin (CFZ, Sigma-Aldrich, St. Louis, MO, USA), cephalothin (CEF, Sigma-Aldrich, St. Louis, MO, USA), cefoxitin (FOX, Sigma-Aldrich, St. Louis, MO, USA), cefuroxime (CXM, Sigma-Aldrich, St. Louis, MO, USA), cefotaxime (CTX, A.G. Scientific, Inc., San Diego, CA, USA), ceftazidime (CAZ, Sigma-Aldrich, St. Louis, MO, USA), ertapenem (ETP, Sigma-Aldrich, St. Louis, MO, USA), imipenem (IPM, Sigma-Aldrich, St. Louis, MO, USA), meropenem (MEM, A.G Scientific, Inc, San Diego, CA, USA) and ampicillin (MEM, A.G Scientific, Inc, San Diego, CA, USA) were performed by broth microdilution in cation-adjusted Mueller-Hinton (MH) broth (Becton Dickinson) with inocula of 5 x 105 CFU/ml in accordance with CLSI MO7-A9 recommendations [27]. All MICs were determined in triplicate at least on three separate occasions to obtain at least 9 discrete data points and compared with EUCAST and CLSI clinical breakpoints for all antibiotics [28, 29]. *E. coli* (ATCC25922) and *Pseudomonas aeruginosa* (ATCC27853) were included in each experiment as quality controls.

### Transfer of resistance genes

The filter mating method [30] was used to transfer plasmids from clinical isolates carrying *bla*_CTX-M-15_, (pJIE143) [31] *bla*_IMP-4_ (pEl1573) [32] and *bla*_KPC-2_ (pJIE2543-1) [33] to *K. pneumoniae* ATCC 13883 and porin mutants (∆OmpK35, ∆OmpK36, OmpK36GD, ∆OmpK35∆OmpK36 and ∆OmpK35OmpK36GD). The presence of resistance genes in transconjugants was confirmed by PCR [33-35] and the presence of plasmids of the expected size confirmed by S1 nuclease pulsed-field gel electrophoresis (Promega, Madison, WI, USA) [36, 37].

### Outer membrane porin investigation

Isolates were grown overnight in MHB. Bacteria were disrupted by sonication and outer membrane porins (OMPs) isolated with sarcosyl (Sigma-Aldrich, St. Louis, MO, USA), as previously described [21, 38]. Samples were boiled, analyzed by sodium dodecyl sulfate-polyacrylamide gel electrophoresis (SDS-PAGE) (12% separating gels), and stained with Imperial protein stain (Thermo Scientific, Rockford, IL, USA), following the manufacturer’s instructions. *K*. *pneumoniae* ATCC 13883, which produces both porins (OmpK35 and OmpK36) was used as a control [39]. Colour prestained protein standard, broad range (11-245 KDa) (New England Biolabs, MA, USA) was used as size marker.

### Real-time reverse transcription-PCR

The expression levels of the different porins were measured by real-time RT-PCR. Cells were harvested in logarithmic phase at an OD_600_ of 0.5-0.6. Total RNA was isolated using RNeasy system (Qiagen). RNA was treated with DNase (TURBO DNA-*free* Kit, Ambion). cDNA was synthesized by high-capacity cDNA reverse transcriptase kit (Applied Biosystems). One microgram of the initially isolated RNA was used in each reverse transcription reaction. cDNA was diluted 1:10 and 2 μl were used for the real-time reaction. Three biological replicates, each with three technical replicates, were used in each of the assays. The relative levels of expression were calculated using the threshold cycle (2^−∆∆*CT*^) method [40]. The expression of *rpoD* was used to normalize the results. The primers used are listed in Table S1.

### Determination of growth rate

Growth rates were determined as previously described [41]. Overnight broth cultures were diluted 1:1000. Six aliquots of 200 μl per dilution were transferred into 96-well microtiter plates (Corning Incorporated, Durham, NC, USA). Samples were incubated at 37°C and shaken before measurement of OD_600_ in a Vmax Kinetic microplate reader (Molecular Devices, Sunnyvale, CA, USA). Growth rates and generation times were calculated on OD_600_ values between 0.02-0.09. The relative growth rate was calculated by dividing the generation time of each mutant by the generation time of the parental strain (*K. pneumoniae* ATCC 13883, 10.85 or 11.76) which was included in every experiment. Experiments were performed in sextuplicate in three independent cultures on three different occasions. Results are expressed as means ± standard errors of the means.

### *In vitro* competition experiment

Competition experiments were carried out as described previously [42]. Viable cell counts were obtained by plating every 24 h on antibiotic-free LB agar and on LB agar supplemented with antibiotic (kanamycin 20 μg/ml or tetracycline 10 μg/ml) to distinguish between mutants and wild-type cells. PCR (with pairs K36GD4 / K36GD11 or K36GD12 / K36GD13 primers, Table S1) was performed for the calculation of the competition results between the parental strain and OmpK36GD mutant (in this particular experiments, bacteria were diluted in fresh media every 24h and PCR on 100 viable colonies of each replicate was performed every 48h). All experiments were carried out in triplicate with three independent cultures. Mean values of three independent experiments ± standard deviation were plotted.

### Mouse model of gastrointestinal tract colonization (GI) and competition experiments

Five to six week-old female BALB/c mice (Animal Resources Centre (ARC), Sydney, Australia) were used for GI colonization [43-45] and competition experiments. Mice were caged in groups of three and had unrestricted access to food and drinking water. Faecal samples were collected and screened for the presence of indigenous *K. pneumoniae* before inoculation. For the colonization study, three mice were inoculated with the parental strain or a porin mutant (1 × 10^10^ CFU / mouse), suspended in 20% (w/v) sucrose. For individual colonization, ampicillin was added to drinking water on day 4 (0.5 g / L) after an inoculation [46]. For the competition experiment, equal volumes of the parental strain and each mutant or equal volume of different mutants (1x 10_10_ CFU / mouse) were mixed and suspended in 20% (w/v) sucrose. Colonization was maintained with ampicillin 0.5 g / L throughout the experiment [47-49]. Faeces samples were collected every second day, emulsified in 0.9% NaCl and appropriate serial dilutions plated on MacConkey-inositol-carbenicillin agar, which selectively recovers for *K. pneumoniae* [50].

### Mouse model of virulence: intranasal infection

Five-six week-old female BALB/c mice [Animal Resources Centre (ARC), Sydney, Australia] used in the inhalation (pneumonia) model [51-53] were exposed to ATCC 13883 and 10.85 and their isogenic ∆OmpK35OmpK36GD mutants. Overnight bacterial cultures were harvested, washed and resuspended at 10^9^ CFU in 20 μl of saline and inoculated into the nasal passages. A control group of mice was inoculated with saline. Following infection, survival studies were performed (10 mice per strain). Organ (lung and spleen) and blood infection burdens were also assessed at various points throughout the infection period, by plating out blood and homogenised tissue onto LB agar, and counting CFU (5 mice per strain, per time point).

### Structure modelling of OmpK36 variants

Tri-dimensional structural models of ATCC13883 OmpK36 and its mutated variant OmpK36GD were computed with ProMod3 Version 1.1.0 on the SWISS-MODEL online server [54] using the target–template alignment method. The best scoring model used as a template was 5nupA (93.84% sequence identity, with a QMEAN equal to −2.29 and −2.14, respectively for both sequences). For comparison purposes, models were also computed using the second best OmpK36 structure available in PDB (1osmA). All predicted models were evaluated using MolProbity [55, 56] and Verify3D [57, 58], with Ramachandran plots generated by MolProbity indicating for all computed models that at least >98% of residues were in allowed regions. Predicted structures were displayed by PyMol software (version 2.1.1) [59].

Additionally, the specific impact of di-nucleotide insertions was also investigated by altering the OmpK6 structure under PDB accession 5nupA, adding either the di-nucleotide GD-, TD- or SD-, after position G113 and modeling the resulting variant sequences in the same manner as mentioned above.

### Genome sequencing and comparative analysis

Genomic DNA was extracted from 2 ml overnight cultures using the DNeasy Blood and Tissue kit (Qiagen). Paired-end multiplex libraries were prepared using the Illumina Nextera kit in accordance with the manufacturer’s instructions. Whole genome sequencing was performed on Illumina NextSeq 500 (150bp paired-end) at the Australian Genome Research Facility (AGRF). Raw sequence reads are available on NCBI under Bioproject accession number PRJNA430457. Reads were quality-checked, trimmed and assembled using the Nullarbor pipeline v.1.20 (available at: https://github.com/tseemann/nullarbor), as previously described [60], but with the exception of the assembly step which was performed using Shovill (available at: https://github.com/tseemann/shovill), a genome assembler pipeline wrapped around SPAdes v.3.9.0 [61] which includes post-assembly correction. Assemblies were also reordered against reference strain *K. pneumoniae* 30660/NJST258_1 (accession number CP006923) using progressive Mauve v.2.4.0 [62] prior to annotation with Prokka [63] and screened for antibiotic resistance genes using Abricate v.0.6 (available at: https://github.com/tseemann/abricate).

### Population analysis

To investigate the significance of *ompK35* and *ompK36* mutations in a wider population, we collected a total of 1,557 draft and complete *K. pneumoniae* genomes publicly available in Genbank (Feb 2017, Table S2). Sequences were typed using Kleborate v0.1.0 [64] to identify MLST (Table S3) and minimum spanning trees were generated using Bionumerics v.7.60. Presence and absence of porins were assessed in the pangenome using Roary v3.6.0 [65] with default parameters and mutations in loop 3 (L3) identified using BLAST. The 2,253,033 nt core genes alignment predicted by Roary was used to build a maximum-likelihood tree using IQ-TREE v1.6.1 [66], with a GTR+G+I nucleotide substitution model and branch supports assessed with ultrafast bootstrap approximation (1000 replicates). Trees were visualized alongside contextual information with Phandango [67].

Statistical analysis was performed using Chi-tests and extended mosaic plots in R, to determine associations between ST, porin defects and other relevant population metrics such as country of origin, year and source of isolation. Relevant R scripts were also made available at https://github.com/nbenzakour/Klebsiella_antibiotics_paper.

### Ethics Statement

For gastrointestinal gut colonization, animal experiments were approved by the Western Sydney Local Health District Animal Ethics Committee (AEC Protocol no. 4205.06.13).

For intranasal infection, animal experiments were approved by the Western Sydney Local Health District Animal Ethics Committee (AEC Protocol no. 4275.06.17).

All research and animal care procedures were in accordance with the “Australian Code of Practice for the Care and Use of Animals for Scientific Purposes”, 8^th^ Edition (2013) (National Health and Medical Research Council, Australian Government).

## Results

### Outer membrane porins and resistance to beta-lactam antibiotics

Minimal inhibitory concentrations for commonly used carbapenems (Ertapenem and Meropenem), third-generation cephalosporins (Ceftazidime, Cefotaxime and Ceftriaxone), cephamycins (also called ‘second generation cephalosporins’, Cefoxitin and Cefuroxime), first generation cephalosporins (Cephalothin and Cefazolin), and the semi-synthetic penicillin Ampicillin were determined in three *K. pneumoniae* strains and their isogenic porin mutants, with representative results in Table 2 (for complete results, see Table S4). The SHV enzyme characteristically expressed by *K. pneumoniae* hydrolyses ampicillin very effectively, providing high MICs to ampicillin [68-71], but does not provide clinically important resistance to cephalosporins or carbapenems in the setting of normal membrane permeability.

Loss of OmpK36 (∆K36 in Tables 2 and S4) is associated with a minor increase in MIC for carbapenems and cephalosporins (Tables 2 and S4), with a lesser impact from OmpK36GD mutations, consistent with an important role for OmpK36 in the nutritious growth media (MHB) normally used for standardised MIC determinations (Tables 2 and S4).

OmpK35 loss (∆K35 in Tables 2 and S4) has little impact alone but further increases MICs for most antibiotics in the presence of OmpK36 lesions (e.g. ∆K35∆K36 and ∆K35∆K36GD). In addition to ertapenem non-susceptibility, ∆OmpK35∆OmpK36 and ∆OmpK35OmpK36GD strains are clinically resistant to first (e.g. cephalothin, CEF) and second generation cephalosporins/ cephamycins (e.g. cefoxitin, FOX) (Tables 2 and S4).

Naturally occurring plasmids from other *K. pneumoniae* strains encoding a common ESBL (*bla*_CTX-M-15_) [31], a metallo-carbapenemase (*bla*_IMP-4_) [32] and a serine-carbapenemase (*bla*_KPC-2_) [33] were transferred into ATCC 13883 and its isogenic mutants by conjugation, with transfer verified by PCR (Table S1) and S1/PFGE (Fig. S1). Even the common ESBL CTX-M-15 confers reduced susceptibility to ETP in the presence of an OmpK36 deletion or inner channel mutation (GD duplication), especially if accompanied by an OmpK35 defect (Table 3). Expression of the specialised carbapenemases IMP and KPC from their naturally occurring plasmids resulted in greatly increased carbapenem MICs (Table 3), with the double porin mutants being highly resistant to all carbapenems tested.

### Altered expression of other common porins in ∆OmpK35, ∆OmpK36, and OmpK36GD

Other porins may compensate for the loss of major outer membrane porins in *K. pneumoniae* [74-77]. Expression of *ompK35*, *ompK36*, *ompK37*, *phoE*, *ompK26* and *lamB* was measured in isogenic porin mutants of ATCC 13883 and 10.85 *K. pneumoniae* strains, in conditions in which either OmpK35 or OmpK36 are ordinarily expressed (Fig 1. Table S5). Neither the introduction of a GD duplication into the OmpK36 inner channel (OmpK36GD) nor the loss of OmpK35 (∆OmpK35 and ∆OmpK35OmpK36GD) affected expression of OmpK36 in MH broth. Loss of OmpK36, however, was associated with increased OmpK35 expression in MH broth, in which OmpK36, but not OmpK35, is ordinarily expressed. Loss of both of these major porins (∆OmpK35∆OmpK36) resulted in increased expression of *phoE* and *lamB*. (Fig 1. Table S5).

**Figure 1:**
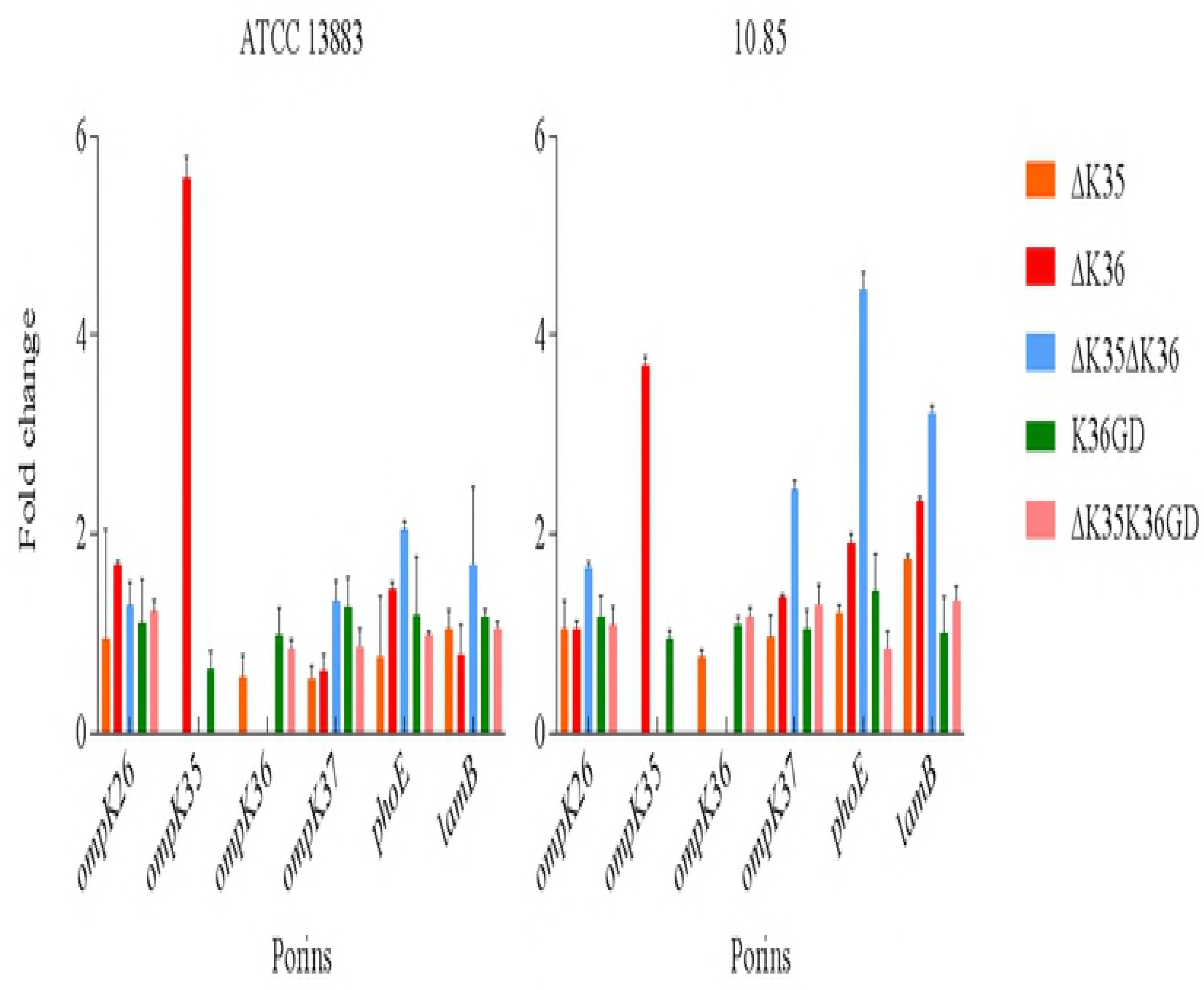
Real-time RT-PCR in *K. pneumoniae* ATCC 13883 and 10.85 porin mutants. The expression of *rpoD* was used to normalize the results. The levels of expression of each mutant are shown relative to the wild type strain ATCC 13883 or 10.85.

### Relative fitness costs of major porin lesions

Exponential phase growth in MH broth was not greatly affected unless both major porins were absent (∆OmpK35∆OmpK36, Table S6) but competition experiments clearly illustrate the importance of OmpK36 (Figs 2 and S2). The ability of ∆OmpK35 strains to directly compete against their intact isogenic parents in MH broth was little affected over 7 days growth (Figs S2A1, S2A3 and S2A4) but OmpK36GD strains are clearly much more able than ∆OmpK36 strains to compete with their isogenic parent strains (Figs 2A1 and 2A3). For ATCC 13883, at day 3, the OmpK36GD population was still 40% of the total combined population (Fig. 2A1), while ∆OmpK36 fell to 20% in the same period (Fig S2B1). This difference was more marked in the presence of an OmpK35 lesion, but ∆OmpK35OmpK36GD populations were still clearly more able than ∆OmpK35∆OmpK36 to compete with the intact parent strain (Figs 2B1, 2B3 versus Figs S2C1 and S2C3). Directly competing OmpK36GD with ∆OmpK36 (and ∆OmpK35OmpK36GD with ∆OmpK35∆OmpK36) further illustrates the competitive advantage, with OmpK36GD strains quickly displacing isogenic ∆OmpK36 strains in MH broth (Figs 2C1 and 2D1). In fact, the introduction of an OmpK36GD mutation had no detectable cost at all in *K. pneumoniae* 10.85 (Figs 2A3 and 2B3), with ∆OmpK35OmpK36GD competing very successfully against the isogenic parent 10.85 (Fig 2B3: 37±4% and 26±15% of the total population represented by ∆OmpK35OmpK36GD on days 6 and 7 respectively)

**Figure 2:**
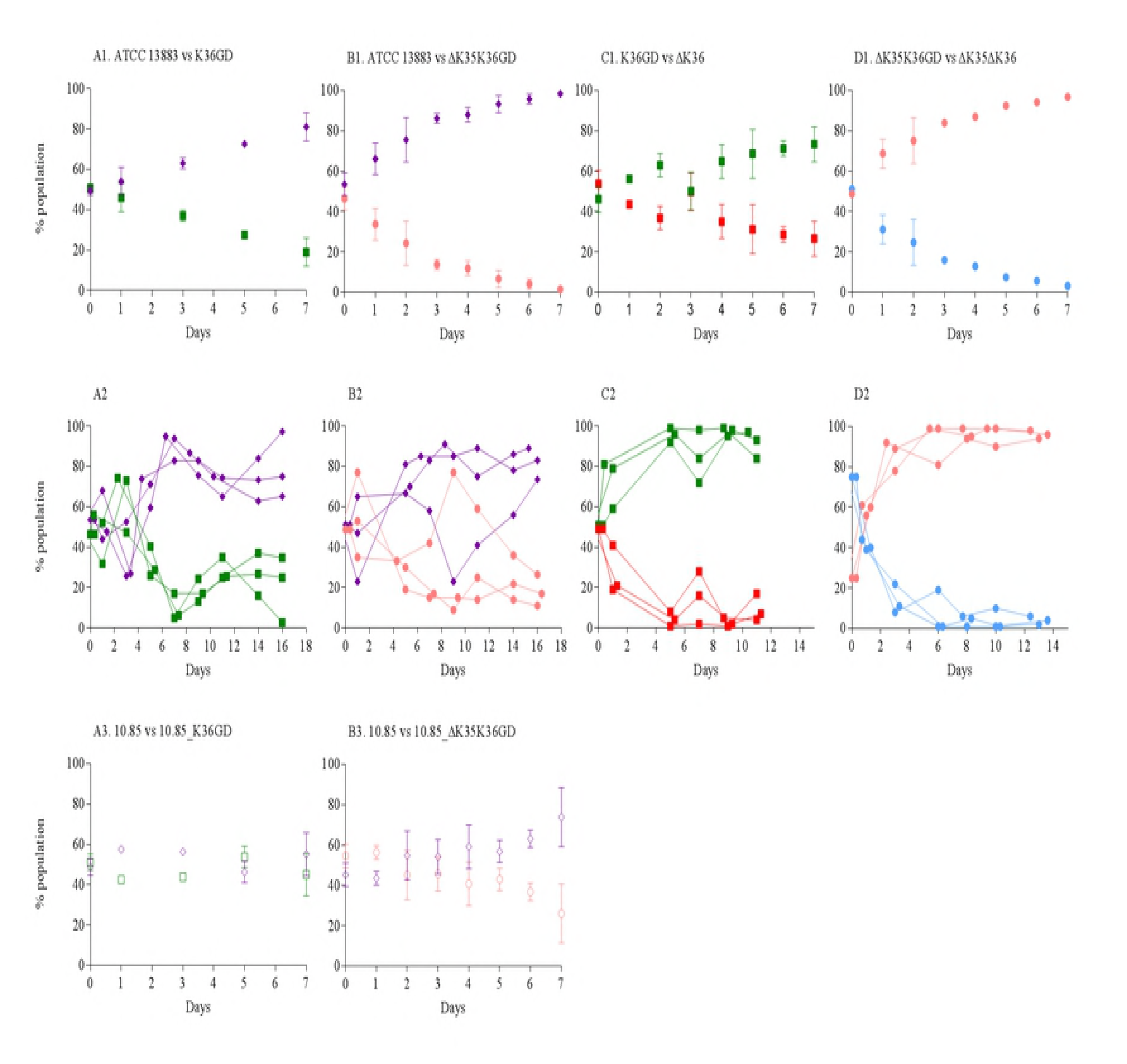
*In vitro* and *in vivo* competition experiments in *K. pneumoniae* ATCC 13883 and 10.85 OmpK36GD porin mutants. The relative fitness of OmpK36GD porin mutants in comparison with parental strain (ATCC 13883 or 10.85) or versus ∆OmpK36 mutant was performed by competition experiments in co-cultures and expressed as a percentage of the mutant or wild type cells versus total population at each time point. *In vitro* growth conditions, MH broth, 37°C. Panels 1 and 3 represent *in vitro* competition experiments for ATCC 13383 and 10.85, respectively. Panels in row 2 show *in vivo* competition experiments for ATCC 13383. For *in vivo* competition experiments, the values for each mouse are represented individually. Violet diamond, ATCC 13883 or 10.85 wild type strains. Green square, OmpK36GD mutant. Pink circle, ∆OmpK35OmpK36GD mutant. Red square, ∆OmpK36 mutant. Blue circle, ∆OmpK35∆OmpK36 mutant.

Mouse gut colonizing studies yielded similar results (Figs 2A2 to 2D2 and S2A2 to S2C2). Mice were confirmed not to include indigenous *K. pneumoniae* on arrival [50], and stable colonisation at ~10^9^ CFU/g faeces was achieved (Fig S3). OmpK35 deficient mutants (∆OmpK35) were not significantly disadvantaged (Fig S2A2) and OmpK36GD strains strongly outperformed OmpK36 strains in competition with their isogenic parents (Figs 2A2 and S2B2). Similarly, direct i*n vivo* competition confirmed a clear fitness advantage of OmpK36GD over ∆OmpK36 (Figs 2C2 and 2D2).

### Pathogenicity is attenuated in ∆OmpK36 but not OmpK36GD or ∆OmpK35 strains

We confirmed the loss of virulence previously ascribed to loss of OmpK36 [78, 79] in a mouse pneumonia model [51-53] and, importantly, showed no difference in lethality between a wild type strain and its isogenic mutant ∆OmpK35/OmpK36GD (Fig 3). Intranasal inoculation of mice showed that these mutations had no significant impact on virulence, with equivalent mortality curves (Figs 3A1 and 3A2) and similar viable counts developing in lung, blood and spleen over the course of infection in isogenic pairs derived from both the ATCC strain and the clinical isolate 10.85 (Figs 3B to 3D).

**Figure 3:**
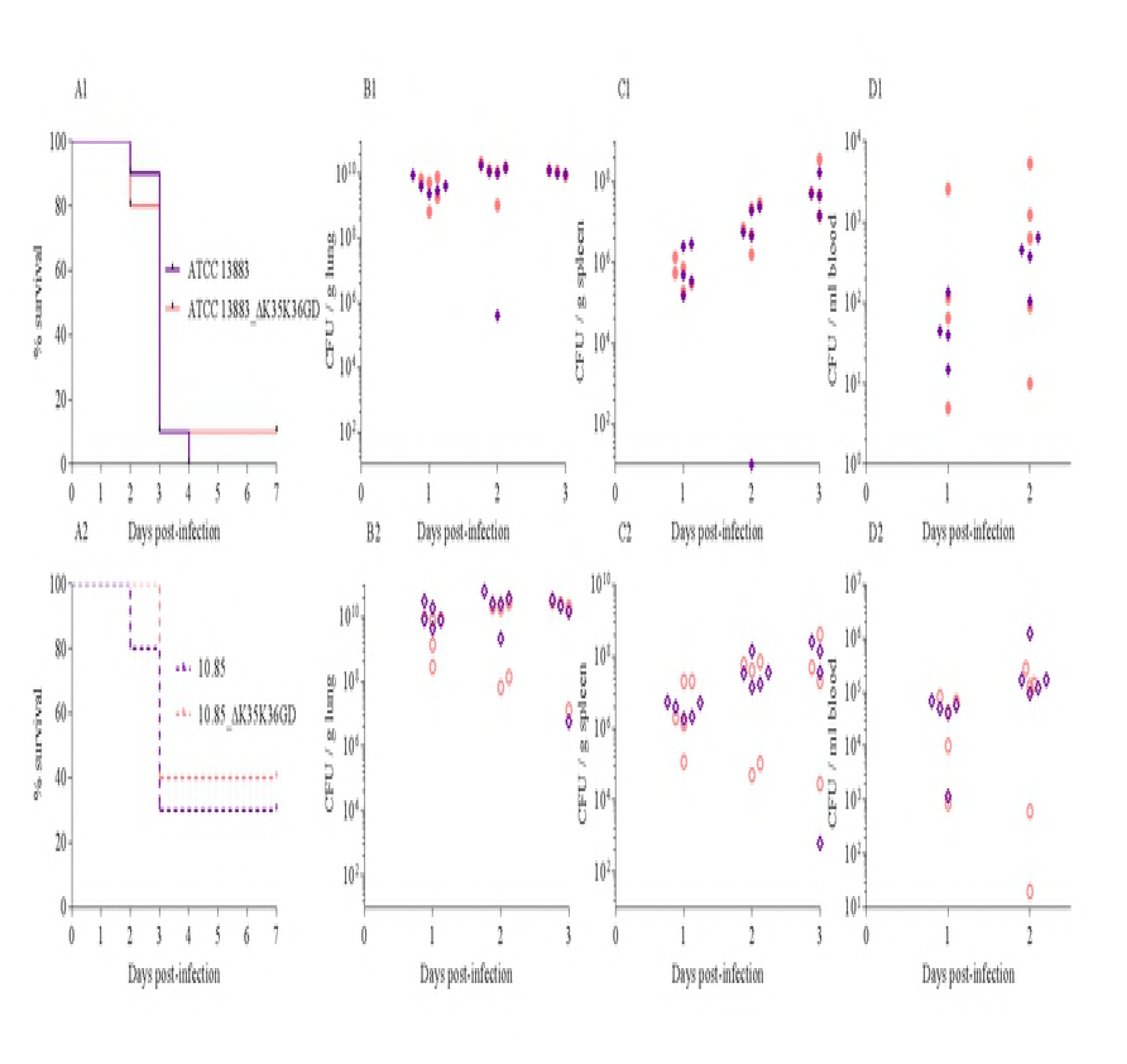
Lung infection experiments in *K. pneumoniae* ATCC 13883 and 10.85 and their isogenic porin mutants ∆OpmK35/OmpK36GD. Survival curves after intranasal infection are represented in panels A1 (for ATCC 13883) and A2 (for 10.85). Organ burden after 24, 48 or 72h is represented in panels B1 to D1 for ATCC 13883 and B2 to D2 for 10.85. Violet diamond, ATCC 13883 or 10.85 wild type strains. Pink circle, ∆OmpK35OmpK36GD mutant.

### Structural impacts of OmpK36 loop L3 mutations

Two crystal structures of native OmpK36 available in the Protein Data Bank under accession number 5nup (2.9 Å, Xray) and 1osm (3.2 Å, Xray) were evaluated as template for structural modelling of OmpK36 and OmpK36GD from ATCC 13883, with targets and templates sharing around 93% nucleotide sequence identity. While Ramachandran plots analysis for all predicted models show at least 98% of residues in allowed regions, other metrics such as QMEAN and Molprobity score are marginally better for ATCC13883 OmpK36 and OmpK36GD models based on the 5nup structure (Fig. 4, Table S7). Although several differences can be observed in the final alignment (Fig 4D), the most prominent differences between the original structure (Fig. 4A) and the ATCC13883 OmpK36 model lie within the loop L6, which can be seen in yellow, slightly obstructing the outmost channel of the porin (Fig. 4B). More strinking is the impact of single di-nucleotide-GD insertion with in loop L3, which is expected to constrict even more the porin channel (Fig. 4C) and is likely responsible for the difference of phenotype between the 2 variants.

**Figure 4:**
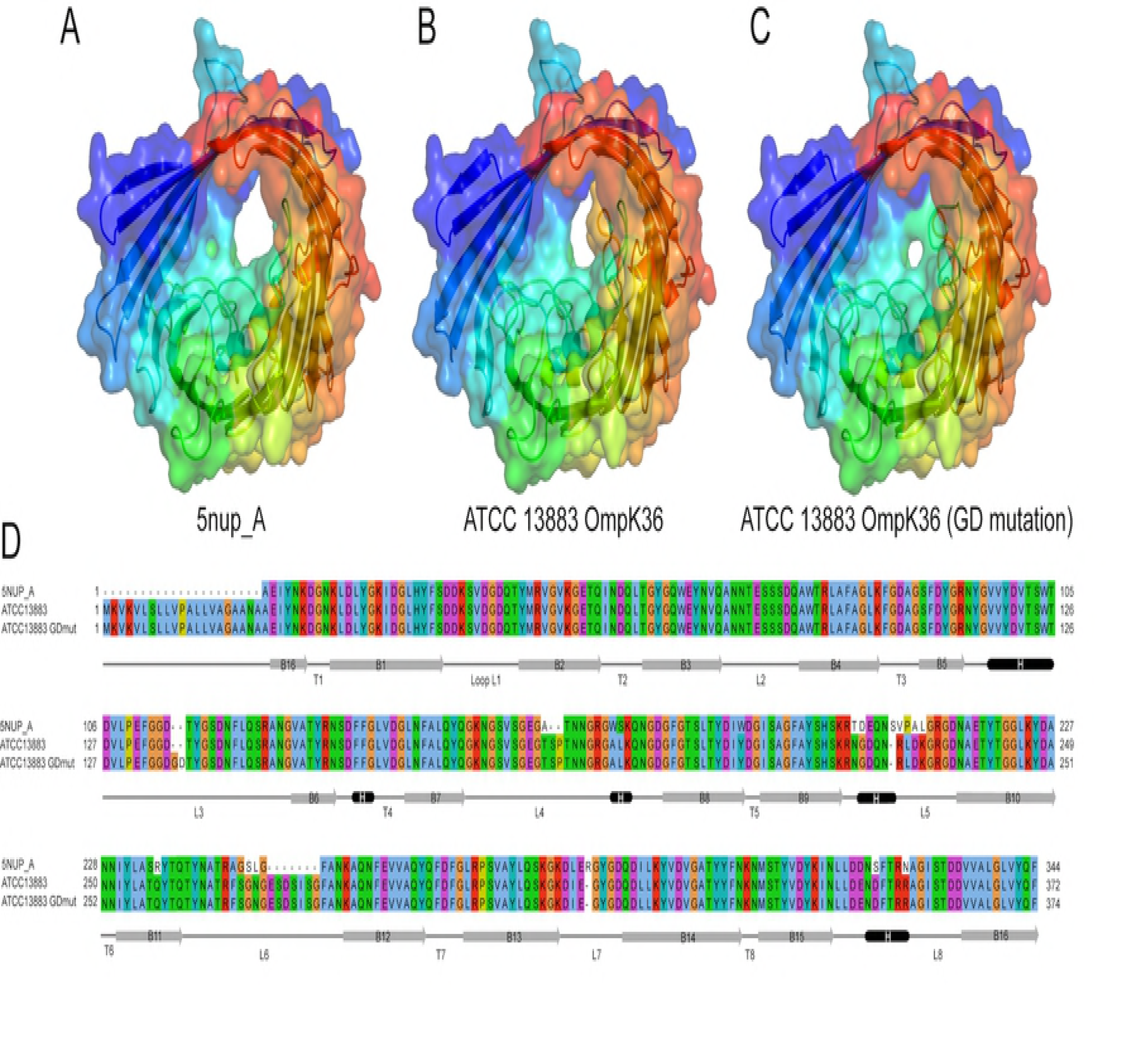
Channel restriction of OmpK36 variants. Comparison of the reference OmpK36 structure under PDB accession 5nupA (A) against predicted structural models of OmpK36 (B) and OmpK36GD mutant (C) from ATCC 13883, showing progressive restriction of the porin channel. The conformation visualised in panel B, in particular the loop 6 in yellow, which can be seen partially obstructing the channel, is not associated with a carbapenem resistance phenotype, contrary to the GD mutant shown in panel C. Panel D consists of the multiple alignment of the 3 corresponding sequences, along with a representation of the predicted secondary structures designated as follows; B for barrel, T for turn, and L for loop. Single peptide is not shown in 5nupA sequence (Panel D).

### OmpK35 loss and convergent evolution of OmpK36GD

The successful antibiotic resistance, colonisation and pathogenicity phenotypes of ∆OmpK35OmpK36GD strains should be reflected in their representation among strains causing human infection. Of 165 unique *K. pneumoniae ompK36* sequences in GenBank, 16% varied from the consensus L3 inner channel motif (PEFGGD). Most common was the GD duplication (PEFGGD**GD**, in 14 of 26 OmpK36 L3 variants identified), along with 6 additional variants: PEFGGD**D**, PEFGGD**SD**, PEFGGD**TD**, PEFGGD**TYD**, PEFGGD**TYG** and PEFGGD**TYGSD** (Fig. S4). Inspection of their corresponding nucleotide sequences suggests that these variants originated from various combinations of short in-frame duplications, combined with additional point mutations in rare cases (Fig. S5).

To investigate OmpK36 among clinical isolates without specialised carbapenemases, we specifically analysed L3 variation in all such *K. pneumoniae* isolates with an Ertapenem MIC > 1 in our local clinical collection (Table 1) by PCR and sequencing (Table S1). Of (n=51), 17 strains (33 %) were identified: all revealed either the previously described GD or TD mutation in the L3 loop of *ompK36* on sequencing and these encoded up to 6 distinct beta-lactamases. These isolates were genetically diverse but belonged to major epidemic clones found elsewhere in the world: *i.e.* ST14, ST16, ST101, ST147, with as many as 6 distinct *ompK35* mutations, all of which introduced disrupting frame-shifts and all of which were relatively lineage-specific (Fig S6).

Finally, all *K. pneumoniae* (complete and draft) genomes available from Genbank, *i.e.* 1,557 entries (as of February 2017) were examined: the two common (GD and TD) variants are shown in a minimum spanning tree built using MLST profiles (Fig. 5) to be distributed across the whole spectrum of diversity of *K. pneumoniae*, including in most major epidemic clones, *e.g.* ST258 and its derivative ST512, ST11, ST101, ST147, ST14 and ST37.

**Figure 5:**
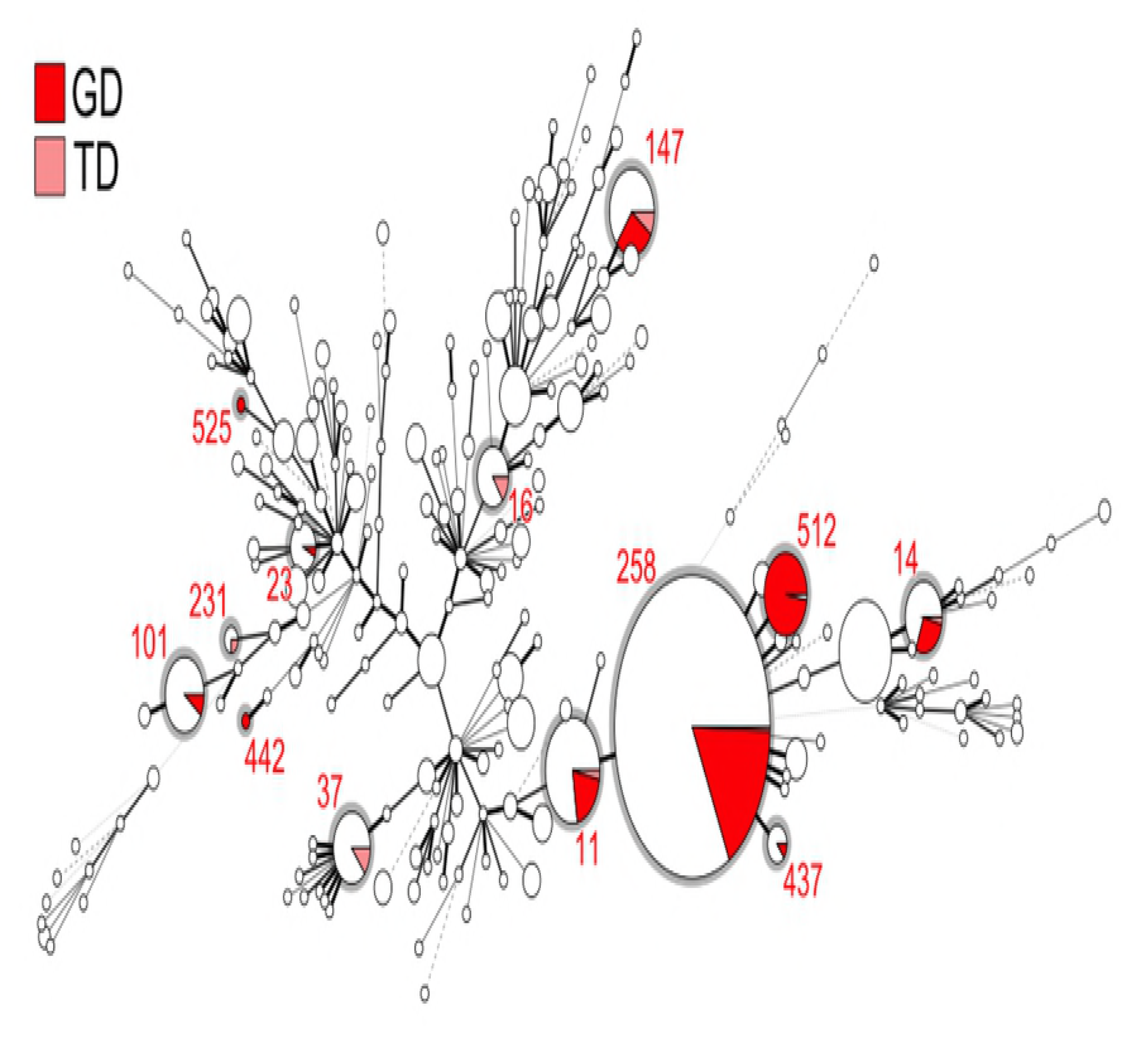
Minimum spanning tree of 1,557 *K. pneumoniae* strains based on their MLST profile. Each circle corresponds to a distinct ST, with its size being proportional to the number of strains of that particular ST (for scale, ST258 contains 552 isolates). Within an ST, the proportion of strains harbouring either a GD or TD insertion in the loop L3 of *ompK36* is shown as a sector coloured in red and pink, respectively. STs carrying these mutations are also circled in grey.

A maximum likelihood phylogeny using a 2,253,033 bp core genome alignment of all 1,557 genomes was computed to contextualize variations in *ompK36* and *ompK35,* with metadata relative to the population (year, source, geographical regions of isolation, as well as major beta-lactamases genes) (Fig. 6). Those genes most relevant to a carbapenem resistance phenotype are shown, and the expected clustering of some of these is as expected (e.g. *bla*_CTX-M-15_ with _OXA-1_ and _TEM-1b_). Major associations with other genes not affected by porin changes are not shown (e.g. aminoglycoside resistance due to 16S methylase genes that are common companions of *bla*_NDM_, other class I integron cassettes from the array in which *bla*_IMP-4_ is found, etc). The predominance of *ompK36* variations in L3 compared to its loss or disruption is evident at a glance, as is the common loss or disruption of *ompK35* in unrelated strains. There is no obvious relationship between OmpK36 L3 variations and the presence of *bla*_KPC_ but there is strong clustering of these variations in certain types (ST 258, 512 etc).

**Figure 6:**
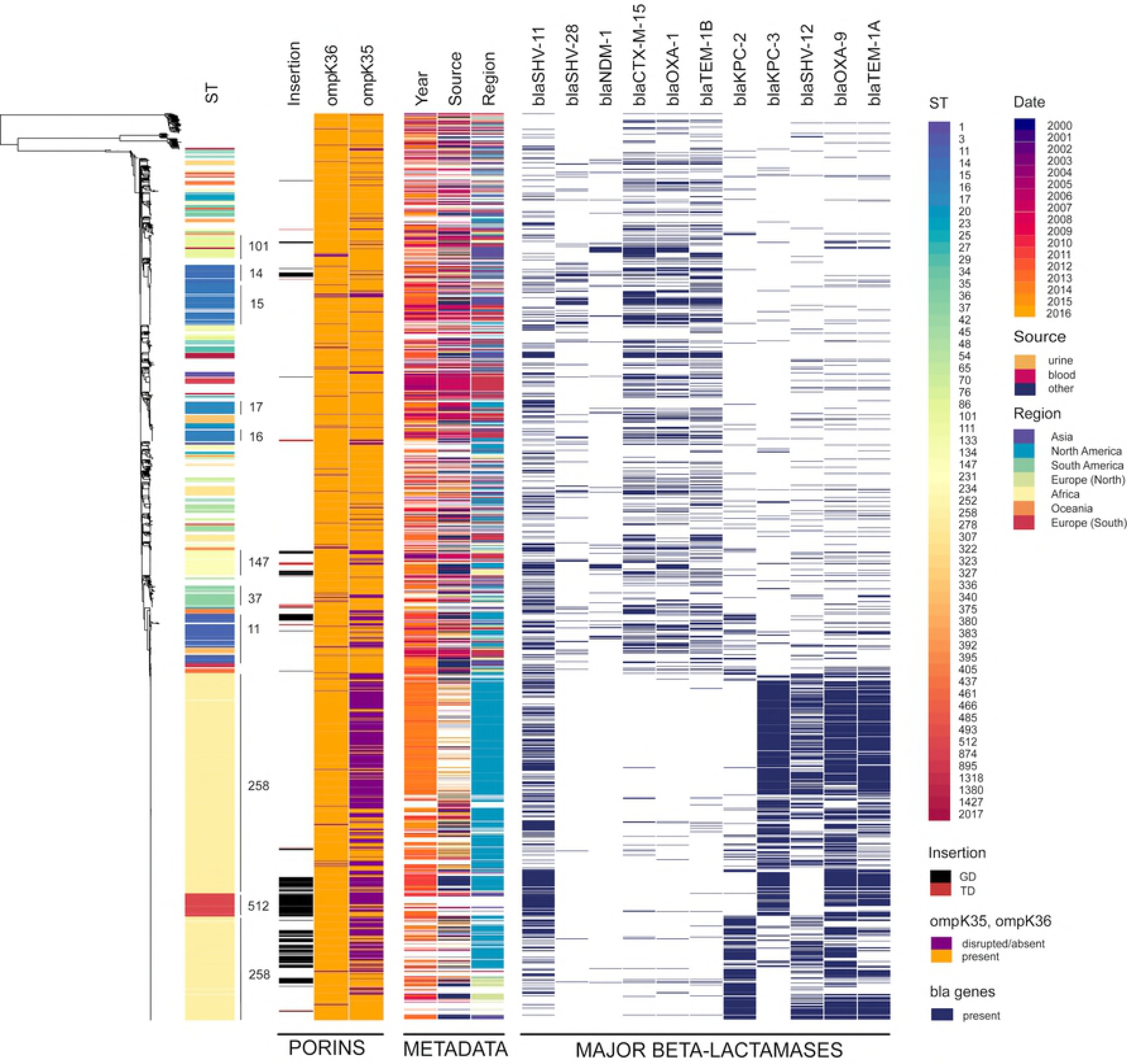
Maximum likelihood tree of 1,557 *K. pneumoniae* strains. A phylogenetic tree was built using a 2,253,033 bp long core alignment. Contextual information relative the collection was visualized using Phandango and includes ST (of which the major ones are indicated on the tree); GD or TD insertion in the loop L3 of *ompK36*, in black and red, respectively; presence or absence of *ompK36*, in orange and purple, respectively; presence or absence of *ompK35*, in orange and purple, respectively. Additional metadata include year of isolation, in a gradient from purple to yellow; source and geographical region of isolation in a rainbow gradient; and presence of major beta-lactamases (bla) alleles identified, in dark blue.

We attempted to take sampling bias into consideration by adjusting for the distribution of samples according to metrics such as ST, country, and year (Fig S7). As expected for a gene so clearly linked to fitness and virulence, *ompK36* is highly conserved across the population (present in 1,499 out of 1,577), independent of ST (Chisq = 207.51, df = 227, p-value = 0.8188), while *ompK35* (evidently dispensable in the host) is disrupted in nearly a third of all strains (Fig. 6), and particularly within the major epidemic clone ST258 (Chisq = 603.7, df = 227, p-value = 5.748e-36). Three-way comparison of the presence/absence of *ompK35*, mutations in *ompK36*, and ST (considering only those STs harbouring *ompK36* GD/TD variants) shows that i) some STs tend to have both *ompK36* and *ompK35* intact (mainly ST15, ST16, ST17); ii) others tend to have intact *ompK36* with disrupted *ompK35* (ST101, ST129, ST258); and iii) some STs tend to have *ompK36* (GD/TD) variants combined with disrupted *ompK35* (ST11, ST14, ST147, ST258 and ST37) (Fig. S8). Associations between presence/absence of OmpK35, an extra aspartate in OmpK36 loop 3 and other metrics such as year or country of isolation may be confounded by over-representation of certain categories (USA, years 2011 and 2014) and we were unable to identify a definite temporal or geographical signal (Fig. S9 and S10).

Finally, we looked at associations between the number of resistance genes and porin defects in major STs, and found that the presence of *ompK36* GD/TD variants did not correlate with a higher number of resistance genes (with the exception of OmpK36GD in ST14). In fact, successful clones such as ST258 and ST11 harbouring OmpK36GD encoded significantly less resistance genes (*p*<0.001, Wilcoxon test) (Fig. S11).

## Discussion

β-lactam antibiotics are among the most commonly prescribed for severe infections [80, 81] and the emergence of β-lactam resistance in *K. pneumoniae* has become a global health threat [82, 83]. In general, *E. coli* and *K. pneumoniae* carrying transmissible β-lactam resistance genes have predictable and normally distributed β-lactam MICs [21] but carbapenem MICs in *K. pneumonia*e are bimodally distributed with higher MICs correlating with OmpK36 defects [21]. OmpK36 loss or mutation is not uncommonly reported in highly resistant clinical isolates producing KPC, ESBL or AmpC β-lactamases [20, 84, 85].

Diffusion of β-lactam antibiotics through non-specific porins such as OmpK35 and OmpK36 is dependent on size, charge and hydrophobicity [86, 87], with bulky negatively charged compounds diffusing at a lower rate than small zwitterions of the same molecular weight [88]. OmpK35 is much less expressed in high osmolarity nutrient-rich conditions than OmpK36, which has the narrower porin channel of the two (Fig S12) [9] and large negatively charged β-lactams such as third-generation cephalosporins and carbapenems diffuse more efficiently through OmpK35 than OmpK36 [78, 89]. Here we confirm the significantly increased MICs, commonly attributed to mutations in these two major porins [10, 90, 91], in three *K. pneumoniae* strains (the widely-published ATCC strain 13883 and two locally isolated clinical strains (Tables 2 and S4); and unequivocally identify the primary role of OmpK36 in carbapenem resistance.

Comparable MIC changes in single (OmpK36GD and ∆OmpK36) and double (∆OmpK35OmpK36GD and ∆OmpK35∆OmpK36) mutants indicate that duplication of a glycine aspartate (GD) pair in a critical position in the porin eyelet region (loop 3) is almost as effective as a complete deletion of the porin in excluding large anionic antibiotics. Both single and double porin mutants were susceptible to extended-spectrum cephalosporins (cefotaxime and ceftazidime) in the absence of acquired hydrolysing enzymes, demonstrating the impotence of the naturally occurring chromosomal SHV [68-70] against these compounds [90].

**Table 2.**
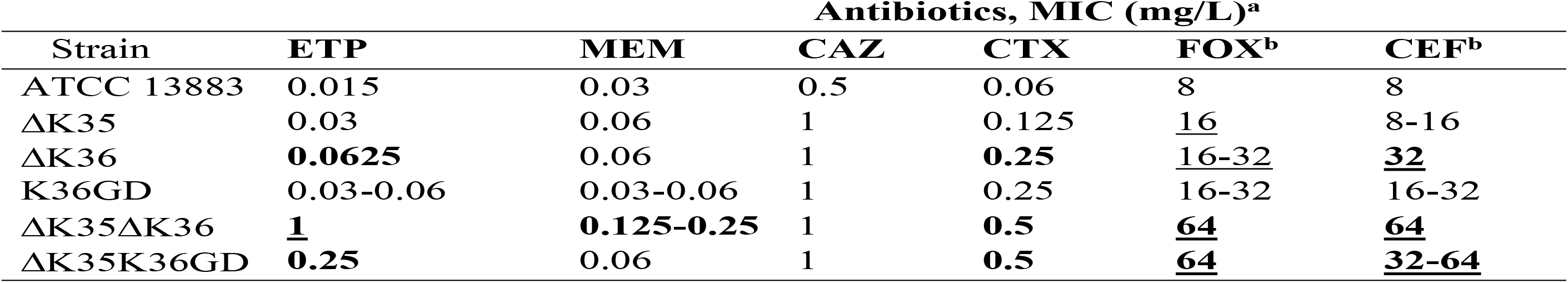
Antibiotic MICs *K. pneumoniae* ATCC 13883 and porin mutants

Differences relating to porin permeability in *K pneumoniae* are most striking and important in the presence of acquired carbapenemases and it is clear that these permeability changes greatly enhance the associated resistance phenotypes. The common Ambler Class A serine protease KPC-2 and Class B metalloenzyme IMP-4 expressed from their natural plasmids produce only borderline resistance against meropenem and the smaller zwitterionic imipenem in the presence of the ‘wild type’ OmpK36 osmoporin (Table 3) but MICs that exceed therapeutic tissue levels [92, 93] are the rule in strains of the commonly occurring ∆OmpK35OmpK36GD genotype.

**Table 3.** Carbapenem MICs against ATCC 13883 and porin mutants with *bla*_CTX-M-15_, *bla*_IMP-4_ or *bla*_KPC_

| Strain | MIC (mg/L) |  |  |  |  |  |  |  |  |
| --- | --- | --- | --- | --- | --- | --- | --- | --- | --- |
|  | CTX-M-15 |  |  | IMP-4 |  |  | KPC |  |  |
|  | ETP | MEM | IPM | ETP | MEM | IPM | ETP | MEM | IPM |
| ATCC 13883 | 0.25 | 0.125 | 1 | <u>8</u> | <u>8</u> | <u>4</u> | <u>16</u> | <u>8</u> | <u>8</u> |
| ΔK35 | 0.5 | 0.125 | 1 | <u>8</u> | <u>8</u> | <u>4</u> | <u>32</u> | <u>32</u> | <u>16</u> |
| ΔK36 | **1** | 0.25 | 1 | <u>8</u> | <u>8</u> | <u>4</u> | <u>32</u> | <u>32</u> | <u>32</u> |
| K36GD | <u>1</u> | 0.25 | 1 | <u>8</u> | <u>8</u> | <u>4</u> | <u>32</u> | <u>16</u> | <u>16</u> |
| ΔK35ΔK36 | **8** | **2** | 1 | **64** | **32** | **64** | **128** | **128** | **128** |
| ΔK35K36GD | **4** | **1** | 1 | **32** | **32** | **16** | **128** | **128** | **64** |

We also show here that the OmpK35 matrix porin has little or no relevance *in vivo* or in test conditions that reliably predict antibiotic efficacy in the clinic (MICs and competitive fitness in Mueller-Hinton broth). Consistent with this, a high percentage of clinical isolates whose genomes have been lodged with GenBank appear to have lost their ability to express OmpK35 altogether (Fig 6). Increased production of the larger channel OmpK35 is expected under low-temperature, low-osmolarity and low nutrient conditions (Fig S12). These favour survival outside the mammalian host and we show that ∆OmpK35 strains fail to compete successfully with their isogenic parents in nutrient-limited conditions (Fig. S13). We confirm that OmpK35 is not naturally expressed at significant levels in optimal growth conditions nor in the mammalian host, as previously described [76, 78] and competition experiments, the most sensitive and direct measures of comparative fitness, evince no discernible disadvantage from the loss of OmpK35 *in vivo,* as expected [19, 94].

Loss of OmpK36 trades off nutrient influx for antibiotic resistance [41, 78] and we show that these more resistant bacteria cannot compete successfully with the antibiotic-susceptible populations from which they arise once antibiotic selection ceases to operate (Fig S2). Double porin mutants (∆OmpK35∆OmpK36) are the most antibiotic-resistant (Tables 2 and S4) but this resistance comes at the cost of a 10% relative growth reduction in nutritious media (Table S6). We show that loss of OmpK36, the main porin normally expressed *in vivo*, is responsible for most of this fitness cost (Figs 2 and S2). The less permeable phosphoporin PhoE and maltodextrin channel LamB are most important in the usual compensatory response when OmpK35 is not available but are not very efficient substitutes (Fig 1 and Table S5). Defects in these porins have been implicated in carbapenem resistance in association with only an AmpC-type enzyme [41, 74, 77, 95] but the fitness cost may be too high for long-term success as such strains are rarely described. By contrast, ∆OmpK35OmpK36GD mutants exhibit little disadvantage compared to isogenic parents with both porins intact, *in vivo* or in optimal growth conditions *in vitro* (Fig 2, Table S6). Expression of OmpK36 is unaffected (Fig 1 and Table S5) as is that of other porins such as OmpK35 (Fig 1 and Table S5), presumably because OmpK36 ‘rescue’ is not required.

The precise loop 3 variation in OmpK36 is evidently a convergent evolutionary process, as a range of different variants occur within genetically distant *K. pneumoniae* populations, all having in common the presentation of an extra negatively charged aspartate (D) residue that significantly constricts the inner channel (Figure 4). The most common solution is the extra glycine and aspartate (PEFGGD to PEFGGDGD in the critical region) which we recreated in isogenic mutants for our experiments. The next most frequent, an extra TD (rather than GD), is similarly likely to spontaneously arise (Fig S5) but is much less common, including in STs in which both GD and TD are found (Figs 5 and 6), implying a less optimal conformation. A recent survey of nearly 500 ertapenem-resistant *Klebsiellae* lacking specialised carbapenemases [96] supports our own finding of the extra aspartate in that position, most commonly as a GD pair, with TD and SD much less often, and other variants being quite rare (Fig. S14). We found no examples of a similarly acidic (glutamate) residue naturally occurring in this position, perhaps reflecting the fact that even simple sequence changes (here, GAY to GAR) add an additional step to a simple duplication event, or the fact that glutamate’s extra carbon makes it slightly less compact than an aspartate in this position.

Other Enterobacteria face the same challenge of excluding bulky anionic carbapenem antibiotics in order to survive high concentrations, even in the presence of a specialist carbapenemase. High level antimicrobial resistance has been ascribed to similar variations in L3 of OmpK36 homologues in *Enterobacter aerogenes*, *Escherichia coli* and *Neisseria gonorrhoeae* (Fig S15) [97-101]. In comparison with their *E. coli* homologues (OmpF and OmpC), OmpK35 and OmpK36 permit greater diffusion of β-lactams [102]. Specifically, OmpK35 appears to be highly permeable to third-generation cephalosporins such as cefotaxime due to its particular L3 domain, which is also seen in Omp35 in *E. aerogenes* but not in other species, and has been proposed as an explanation for the high proportion of *K. pneumoniae* clinical isolates that lack this porin [102, 103]. Our findings of increased MICs in OmpK35 mutants are consistent with those of others [102] but we show here that the more permeable OmpK35 is not important in the mammalian host. Rather, the much less permeable OmpK36 (equivalent to *E coli* OmpC) [102] is the bottleneck for large anionic antibiotics.

The term ‘high risk clone’ [104, 105] is given to host-adapted/ pathogenic strains that dominate the epidemiology of (antibiotic resistant) infections, presumably because they are more transmissible, more pathogenic and/or more tolerant of host-associated stresses (including antibiotics). Here, we see a range of unrelated clonal groups already identifiable as high-risk clones that are dispensing with the OmpK35 porin (Fig 6). The minimal antibiotic resistance advantage in nutritious media is only evident with carbapenems and is unlikely to arise in the presence of an existing OmpK36 loss mutation because the fitness cost is substantial. The loss of OmpK35 through low-level carbapenem exposure in environmental conditions is possible [106] but also has a marked fitness cost and the exposure to carbapenems in the environment is expected to be limited, as they are a still a minority class of prescribed antibiotics and are not yet as common in environmental waters as the sulfonamides, quinolones, macrolides, tetracyclines and other beta-lactams [107].

A recent review of antibiotic resistance in *Klebsiella* pointed out that “The exact role of porins in antimicrobial resistance is difficult to determine because other mechanisms…are commonly present…” [108]. We suggest that host-adaptation in *K. pneumoniae* is widespread and that many *K. pneumoniae* have dispensed with the OmpK35 matrix porin required for an environmental life cycle. Under a major stress (as antibiotic pressure or high concentrations of bile salts in the intestinal lumen), bacteria try to adapt to the new environment [109]. It has been described that for *E. coli*, toxic agents as antibiotics and bile acids diffuse better through the larger OmpF channel (homolog of OmpK35) than the narrower OmpC (equivalent to OmpK36 in *K. pneumoniae*) [110]. At the same time, high osmolarity, high temperature, low pH and anaerobiosis (typical conditions in gut environment) induce the production of OmpK36 but inhibit the expression of *ompK35* [111] [112] [113]. Interestingly, *E. coli* mutants with reduced permeability (decreased *ompF* and increased *ompC* mRNA and protein levels compared with parental strain) can be easily recovered from intestinal gut of germ-free mice after few days of colonization [114]. Convergent evolution upon a highly specific variation in the inner channel of the osmoporin OmpK36 efficiently solves the problem of carbapenem resistance at no cost to colonising ability, competitiveness or pathogenicity and can be expected to be an increasingly common feature of host-adapted ‘high-risk clones’.

There are three direct and immediate implications. Firstly, efforts to control the spread of such strains will be facilitated to some extent by the loss of environmental hardiness resulting from OmpK35 deletion, and should shift slightly more toward managing interpersonal transmission. Secondly, *K. pneumoniae* can be expected to become more antibiotic resistant overall and organisms expressing currently circulating plasmid-borne carbapenemases will more commonly be untreatable with carbapenem antibiotics (e.g. ST258 strains with *bla*_KPC_); the second (higher MIC) peak in the bimodal distribution of carbapenem MICs in *K. pneumoniae* populations will become more prominent. Finally, the mobile carbapenemase gene pool can be expected to flourish in the protected niche provided by host-adapted *K. pneumoniae* populations under strong carbapenem selection pressure in human hosts, thereby increasing the general availability of highly transmissible carbapenem resistance plasmids among host-adapted pathogens in the *Enterobacteriaceae*.

## Supplemental information

**Figure S1: S1/PFGE of strains with the conjugated plasmids.** White arrows show the plasmids in original host isolates and transconjugants. MW; Mid-range PFG Marker.

**Figure S2: *In vitro* and *in vivo* competition experiments in *K. pneumoniae* ATCC 13883, 10.85 and 11.76 knock-out porin mutants.** The relative fitness of deletion porin mutants in comparison with parental strain (ATCC 13883, 10.85 or 11.76) was performed by competition experiments in co-cultures and expressed as a percentage of the mutant or wild type cells versus total population at each time point. *In vitro* growth conditions, MH broth, 37oC. Panels 1, 3 and 4 represent *in vitro* competition experiments for ATCC 13383, 10.85 and 11.76, respectively. Panels in row 2 show *in vivo* competition experiments for ATCC 13383. For *in vivo* competition experiments, the values for each mouse are represented individually. Violet diamond, ATCC 13883, 10.85 or 11.76 wild type strains. Orange square, ∆OmpK35 mutant. Red square, ∆OmpK36 mutant. Blue circle, ∆OmpK35∆OmpK36 mutant.

**Figure S3: Individual gut colonization.** *K. pneumoniae* intestinal colonization in a mouse model. CFU counts of *K. pneumoniae* ATCC 13883 and porin mutants from mice faecal sample. Bacterial inoculum at day 0 is 1×1010 CFU /mouse. Addition of ampicillin 0.5 g / L in the drinking water on day 4. Violet diamond, ATCC 13883 wild type strains. Orange square, ∆OmpK35 mutant. Red square, ∆OmpK36 mutant. Blue circle, ∆OmpK35∆OmpK36 mutant. Green square, OmpK36GD mutant. Pink circle, ∆OmpK35OmpK36GD mutant.

**Fig S4: Alignment of *Klebsiella pneumoniae* OmpK36 proteins**. 26 unique sequences with OmpK36 L3 variants from GenBank were compared with OmpK36 of NTUH_K2044. Isolates with wild-type L3 sequence are not included. Dot line, signal peptide. Black line, beta strands. Red line, loops. Blue line, alpha helix. Green squares, turns. OmpK36 secondary structure based on previous studies. (1, 2). Red boxes, residues involved in the pore eyelet based on (3). Black box, L3 variants.

1. Cowan SW, Schirmer T, Rummel G, Steiert M, Ghosh R, Pauptit RA, et al. Crystal structures explain functional properties of two *E. coli* porins. Nature. 1992;358(6389):727-33.

2. Domenech-Sanchez A, Hernandez-Alles S, Martinez-Martinez L, Benedi VJ, Alberti S. Identification and characterization of a new porin gene of *Klebsiella pneumoniae*: its role in β-lactam antibiotic resistance. J Bacteriol. 1999;181(9):2726-32.

3. Bornet C, Saint N, Fetnaci L, Dupont M, Davin-Regli A, Bollet C, et al. Omp35, a new Enterobacter aerogenes porin involved in selective susceptibility to cephalosporins. Antimicrob Agents Chemother. 2004;48(6):2153-8.

**Figure S5: Proposed scenario of the major duplications observed in OmpK36 loop 3.** Based on observations of the codon sequences, the extra –SD and –SYG following GGD likely result from a combination of duplication followed by point mutation.

**Figure S6: Phylogenetic tree of an Australian collection of *K. pneumoniae* isolates with various degrees of non-susceptibility to carbapanems.** Metadata includes year of isolation; MIC levels for ETP: ertapenem, IMP: imipenem, and MEM: meropenem; *ompK36* L3 mutation; *ompK35* disrupted mutations (as listed in Table S7); ST: sequence type; number of predicted resistance genes encoded; carbapenamase gene encoded; ESBL: extended-spectrum beta-lactamase gene encoded.

**Figure S7: Imbalanced features of the 1,557 *Klebsiella pneumoniae* genomes found in Genbank** (February 2017). Bubble chart showing the distribution of isolates across countries with at least 2 *Klebsiella pneumoniae* genomes reported, from 2001 to 2016, and coloured according to their MLST. The size of each circle is proportional to the number of isolates.

**Figure S8: Extended mosaic plot of the observed proportions of isolates with porins OmpK35 and OmpK36 variations, across STs harbouring *ompK36* GD/TD variants.** The mosaic plot shows the relationships between 3 variables; ST (in purple) and presence/absence of *ompK35* (in black) on the *x*-axis; and presence/absence and mutations of *ompK36* (in grey) on the *y*-axis. The size of each plot tile is proportional to counts. Plot tiles are colored according to their standardized Pearson residuals, as determined by a log-linear model. Deeper shades of red and blue corresponding to a standardized residual less than-4 or greater than +4, respectively, can be interpreted as combinations observed significantly less or more than expected (under the assumptions that proportions have equal levels).

**Figure S9: Extended mosaic plot of the observed proportions of isolates with porins OmpK35 and OmpK36 variations versus years.** The mosaic plots show the relationships between 2 variables; A) year of isolation on the *x-*axis and presence/absence of *ompK35* on the *y*-axis; B) year of isolation on the *x*-axis, and presence/absence and mutations of *ompK36* on the *y*-axis. The size of each plot tile is proportional to counts. Plot tiles are colored according to their standardized Pearson residuals, as determined by a log-linear model. Deeper shades of red and blue corresponding to a standardized residual less than-4 or greater than +4, respectively, can be interpreted as combinations observed significantly less or more than expected (under the assumptions that proportions have equal levels).

**Figure S10: Extended mosaic plot of the observed proportions of isolates with porins OmpK35 and OmpK36 variations versus countries**. The mosaic plots show the relationships between 2 variables; A) country of isolation on the *x-*axis and presence/absence of *ompK35* on the *y*-axis; B) country of isolation on the *x*-axis, and presence/absence and mutations of *ompK36* on the *y*-axis. The size of each plot tile is proportional to counts. Plot tiles are colored according to their standardized Pearson residuals, as determined by a log-linear model. Deeper shades of red and blue corresponding to a standardized residual less than-4 or greater than +4, respectively, can be interpreted as combinations observed significantly less or more than expected (under the assumptions that proportions have equal levels).

**Figure S11: Distribution of resistance genes identified in ST harbouring OmpK36GD or OmpK36TD mutants.** Boxplots were used to display the distribution of resistance genes identified with Abricate within each ST with the following OmpK36 variants, namely isolates with –GD in bright red, –TD in brown, or no insertion (–) in grey. Mean comparison *p*-values are also shown for each ST (Wilcoxon test, with ‘-’ used as a reference group; ns: *p* > 0.05; *: *p* <= 0.05; **: *p* <= 0.01; ***: *p* <= 0.001; ****: *p* <= 0.0001). In addition, the corresponding underlying isolate population is also visualised as individual points, coloured according to OmpK35 type, (1) intact in turquoise or (0) disrupted in coral.

**Figure S12: SDS-PAGE analysis of outer membrane porins.** Wild type strains ATCC 13883, 10.85 and 11.76 were cultured under different temperatures (37°C, 30°C and 25°C) and different nutrient concentrations (MH and MH 1:10). Blue arrow, OmpK35. Black arrow, OmpK36. Red arrow, OmpA.

**Figure S13: *In vitro* competition experiments in MH 1:10 dilution**. The relative fitness of porin mutants in comparison with parental strain ATCC 13883 was performed by competition experiments in co-cultures and expressed as a percentage of the mutant or wild type cells versus total population at each time point. *In vitro* growth conditions: A, MH 1:10 broth, 25°C; B, MH 1:10 broth, 37°C. Violet diamond, ATCC 13883. Orange square, ∆OmpK35. Red square, ∆OmpK36.

**Figure S14: Distinct channel restrictions of OmpK36 di-nucleotide mutants (–GD, –TD, and –SD).** Comparison of the reference OmpK36 structure under PDB accession 5nup1A (WT, wild type) against predicted structural models of mutants harbouring a di-nucleotide insertion in loop 3 after G113, namely GGDGD, GGDTD and GGDSD. For each predicted structure, the 2 most protruding amino-acids resulting from the di-nucleotide insertion were marked and coloured according to their backbone structure (carbons in yellow, oxygens in red and nitrogens in blue).

**Figure S15**: **A. Alignment of *E. coli* OmpC_L3 variants**. 11 unique Omp36_L3 variants from GenBank were compared with L3 of OmpC of K-12 MG1655 (NP_416719). Black boxes, residues different from NP_416719. **B. Alignment of *E. aerogenes* Omp36_L3 variants**. Four unique Omp36_L3 variants from GenBank were compared with L3 of Omp36 from ATCC 13048 (AF335467). Black boxes, residues different from AF335467.

Isolates with wild-type L3 sequence are not included. Black line, loop 3. OmpK36 L3 location based on previous studies. (1, 2). Red boxes, residues involved in the pore eyelet based on (3).

1. Cowan SW, Schirmer T, Rummel G, Steiert M, Ghosh R, Pauptit RA, et al. Crystal structures explain functional properties of two *E. coli* porins. Nature. 1992;358(6389):727-33.

2. Domenech-Sanchez A, Hernandez-Alles S, Martinez-Martinez L, Benedi VJ, Alberti S. Identification and characterization of a new porin gene of *Klebsiella pneumoniae*: its role in β-lactam antibiotic resistance. J Bacteriol. 1999;181(9):2726-32.

3. Bornet C, Saint N, Fetnaci L, Dupont M, Davin-Regli A, Bollet C, et al. Omp35, a new Enterobacter aerogenes porin involved in selective susceptibility to cephalosporins. Antimicrob Agents Chemother. 2004;48(6):2153-8.

**Table S1: Primers and plasmids used during this work.**

**Table S2: Genbank metadata.**

**Table S3: Resistance and typing screening.**

**Table S4: Antibiotic MICs against *K. pneumoniae* and porin mutants.**

**Table S5: Real-time RT-PCR in *K. pneumoniae* ATCC 13883 and 10.85 porin mutants.** The expression of *rpoD* was used to normalize the results. The levels of expression of each mutant are shown relative to the wild type strain ATCC 13883 or 10.85.

**Table S6: Relative growth rate and doubling time of *K. pneumoniae* and porin mutants.**

**Table S7: Model evaluation results ATCC 13883 OmpK36 L3variants.**

